# Entropic interfacial charge separation suppresses coarsening of nanoscale condensates

**DOI:** 10.1101/2023.10.06.561146

**Authors:** Feipeng Chen, Jiaxing Yuan, Yaojun Zhang, Hajime Tanaka, Ho Cheung Shum

## Abstract

Droplet coarsening is a long-standing phenomenon widely observed in our daily life and industrial processes. This process is typically governed by classic theories, such as Brownian motion-induced coalescence and Ostwald ripening, predicting continuous and rapid droplet growth. However, recent studies revealed that nanoscale biomolecular condensates, formed by liquid-liquid phase separation (LLPS), often defy this expectation, exhibiting remarkable long-term stability in cells and in vitro systems. Here, we reveal a merging-limited coarsening mechanism that underpins this anomalously slow growth. Using experiments, theory, and simulations, we demonstrate that nanoscale coacervates formed at neutral stoichiometry remain stable over extended periods due to size-dependent merging inefficiency. This inefficiency stems from entropic charge separation caused by asymmetric chain lengths of oppositely charged polymers, which induces interfacial charge accumulation and inter-coacervate electrostatic repulsion. Our findings reframe LLPS as a kinetically constrained process evolving over a rugged energy landscape, in which merging barriers trap condensates in metastable, long-lived states. This framework offers a physical basis for condensate size control in cells and a design principle for stable synthetic biomolecular assemblies.

## Introduction

Droplet coarsening occurs across a wide range of natural and industrial processes, from oil droplets merging in stirred salad dressing to water droplets condensing on windows. Coarsening is a spontaneous process in which droplets merge and ripen over time, driven by the system’s tendency to minimize interfacial free energy and approach thermodynamic equilibrium. Conventionally, droplet coarsening proceeds through two main mechanisms: Brownian motion-induced coalescence (BMC) ^1–3^ and Ostwald ripening driven by pressure gradients ^4,5^. There has been a growing emphasis on understanding analogous coarsening phenomena in living cells, prompted by the recent discovery of biomolecular condensates and their critical roles in cellular physiology and diseases ^1,4,6–8^; in particular, many condensates exhibit liquid-like behaviors including fusing upon contact, wetting surfaces, and dripping under shear stresses ^9–11^.

Unlike classic coarsening, many condensates in living cells maintain stable sizes and rarely grow over long time periods ^1,4,12–14^. This stability is often attributed to intracellular complexity, where various factors influence coarsening. For example, active reactions can regulate droplet size ^15–18^, whereas nuclear chromatin and cytoplasmic actin passively suppress BMC and Ostwald ripening of condensates ^1,4,13^. In addition, absorption of protein-RNA clusters onto condensate surfaces can form “Pickering emulsions”, wherein the absorbed clusters act like surfactants to reduce the surface tension and create physical barriers to merging, collectively impeding the coarsening process ^12,19^. However, these mechanisms depend on specific cellular conditions to suppress the droplet coarsening.

Therefore, in vitro condensates, which lack such cellular features, are generally expected to coarsen rapidly via classical mechanisms. Indeed, this has been observed for most micrometer-sized condensates in previous studies (Fig. 1a) ^2^. However, recent experiments challenge this expectation. Surprisingly, nanoscale condensates comprised of proteins remain stable over extended periods even in dilute, simple aqueous solutions (Fig. 1a) ^14,20–24^. This contradicts with predictions of BMC and Ostwald ripening ^25,26^, raising a fundamental question: what stabilize these nanoscale condensates?

**Fig. 1.**
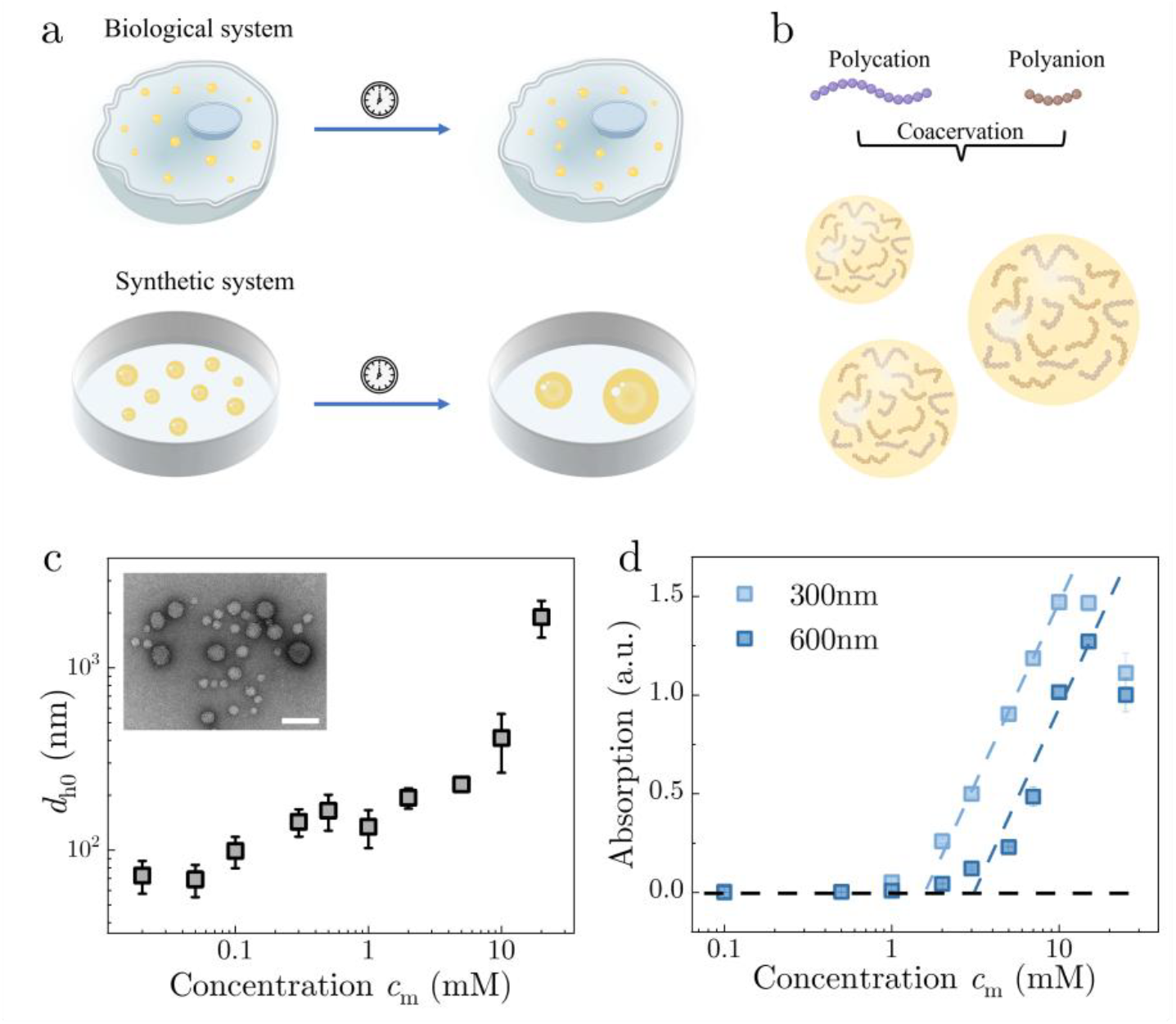
Complex coacervates formed by mixing oppositely charged polyelectrolytes with asymmetric chain lengths. (a) Schematic illustration of distinct coarsening dynamics of condensates in cells and synthetic systems. In cells, condensates remain stable over time while they often merge and grow into larger ones quickly in synthetic systems. (b) Schematic illustration of complex coacervates formed by mixing oppositely charged polycations and polyanions with asymmetric chain lengths at a charge-balanced stoichiometry ratio. (c) The initial mean hydrodynamic diameter *d*_h0_ measured immediately after mixing as a function of monomer concentration of polyelectrolyte *c*_m_. The insert TEM image shows nanoscale coacervates formed at *c*_m_ = 0.05 mM. The white scale bar is 100 nm. (d) The absorption values of coacervate solutions as a function of monomer concentration, measured at two distinct wavelengths, 300 nm and 600 nm. Error bars represent mean ± standard deviation (SD) calculated from at least three independent samples.

In this work, we combine experiments, theory, and simulations to reveal a general merging-limited coarsening (MLC) mechanism that governs the long-time stability of nanoscale condensates. We show that coacervates formed from oppositely charged polyelectrolytes at neutral stoichiometry (i.e., equal charge balance by monomer concentration, not chain number) exhibit suppressed growth at the nanoscale, while larger ones follow classical BMC. Notably, merging efficiency decreases sharply with droplet size, becoming extremely low below a few hundred nanometers. To explain this, we develop a theoretical model that incorporates size-dependent merging probability, supported by coarse-grained simulations. We find that chain-length asymmetry induces entropic charge separation, leading to interfacial charge buildup and electrostatic repulsion that suppress merging. These findings challenge the classical view of LLPS as a purely thermodynamically driven process. Instead, we show that kinetic constraints—arising from molecular-scale features such as chain-length asymmetry and interfacial charge separation— impose size-dependent electrostatic barriers that arrest nanoscale condensates in metastable states. As a result, LLPS in such systems proceeds not along a smooth thermodynamic descent but over a rugged energy landscape shaped by interfacial physics and molecular interactions. This work thus reframes LLPS as a kinetically constrained phenomenon, governed by electrostatic merging barriers and entropic partitioning, offering a new physical framework for understanding and controlling condensate stability in biological and synthetic systems.

## Results

### Formation of nanoscale complex coacervates at low polyelectrolyte concentrations

Many biological condensates form through the complex coacervation between proteins and nucleic acids via multivalent interactions ^11,27^. To understand the physical principles underlying biological phase separation, complex coacervates comprised of oppositely charged polyelectrolytes have been widely employed as model systems due to their simplified physicochemical properties ^28–30^. In particular, complex coacervates resemble representative characteristics of biological condensates, such as selective encapsulations of molecules ^31^, multiphase organizations ^32^, enhancement of biochemical reactions ^33^, and aging kinetics ^34^.

In this work, we investigate the coarsening dynamics of complex coacervates across different size scales. These coacervates are formed by mixing solutions of negatively charged poly(methacrylic acid, sodium salt) (PMA) and positively charged poly(diallyldimethylammonium chloride) (PDDA) at a 1:1 stoichiometric ratio of monomer concentrations (Fig. 1b). Using dynamic light scattering (DLS), we measure the mean hydrodynamic diameters (*d*_h0_) of coacervates formed immediately after mixing. From the experimental results, *d*_h0_ increases monotonically with increasing monomer concentration (*c*_m_) (Fig. 1c). Additionally, turbidity measurements of coacervate solutions at different monomer concentrations show a rapid increase in turbidity once a threshold is crossed followed by a plateau regime (Fig. 1d).

To confirm the DLS results, coacervates are observed using an optical microscope. At a high concentration of *c*_m_ = 20 mM, micrometer-sized coacervates are clearly observed (Supplementary Fig. 1d). At intermediate concentrations (*c*_m_ = 0.5 mM and 2 mM), nanoscale coacervates are observed as blurry shadows in bright-field images (Supplementary Fig. 1b). At even lower concentrations, nanoscale coacervates (i.e., sizes below a few hundred nanometers) are not discernible under optical microscopy (Supplementary Fig. 1a). These nanoscale condensates are further visualized clearly using transmission electron microscopy (TEM) (Supplementary Fig. 2). In magnified images, polymer brushes at the periphery and a condensed core are identified, indicating a gradual density gradient between coacervates and the surrounding solution (Supplementary Fig. 2d). Notably, coacervate diameters measured from TEM images are comparable to *d*_h0_ obtained by DLS.

### Suppressed coarsening dynamics of nanoscale coacervates

The above results are consistent with recent studies showing the formation of nanoscale clusters at low concentrations ^23,25,35,36^. However, how these nanoscale droplets evolve over time has not been thoroughly explored. Here, we further use DLS to monitor the evolution of hydrodynamic diameters *d*_*h*_ of coacervates formed at various concentrations over a long-time duration, typically exceeding 12 hours. Phase separation generally initiates with a nucleation or spinodal decomposition stage, followed by a coarsening process in later phases ^2,3^. Given that the initial stage is typically fast ^20,37,38^, our measurements primarily capture the later coarsening stage. DLS measurements reveal that microscale coacervates formed at *c*_m_ = 16 mM exhibit a power law growth of *d*_*h*_ with an exponent close to 1/3 (Fig. 2a). This coarsening behavior aligns with previous studies and is well described by classic theories, such as Brownian motion-induced coalescence (BMC) or Ostwald ripening ^2,3^. Surprisingly, at intermediate concentrations (*c*_m_ = 2 mM and 5 mM), nanoscale coacervates (~200 nm) initially remain stable, yet undergo abrupt, rapid growth in later stages, with power-law exponents deviating from 1/3 (Fig. 2b). More intriguingly, nanoscale coacervates formed at *c*_m_ = 0.1 mM and *c*_m_ = 0.3 mM remain stable over a period of 12 hours with their hydrodynamic diameters *d*_h_ remaining at ~80 nm and ~150 nm, respectively (Fig. 2c).

**Fig. 2.**
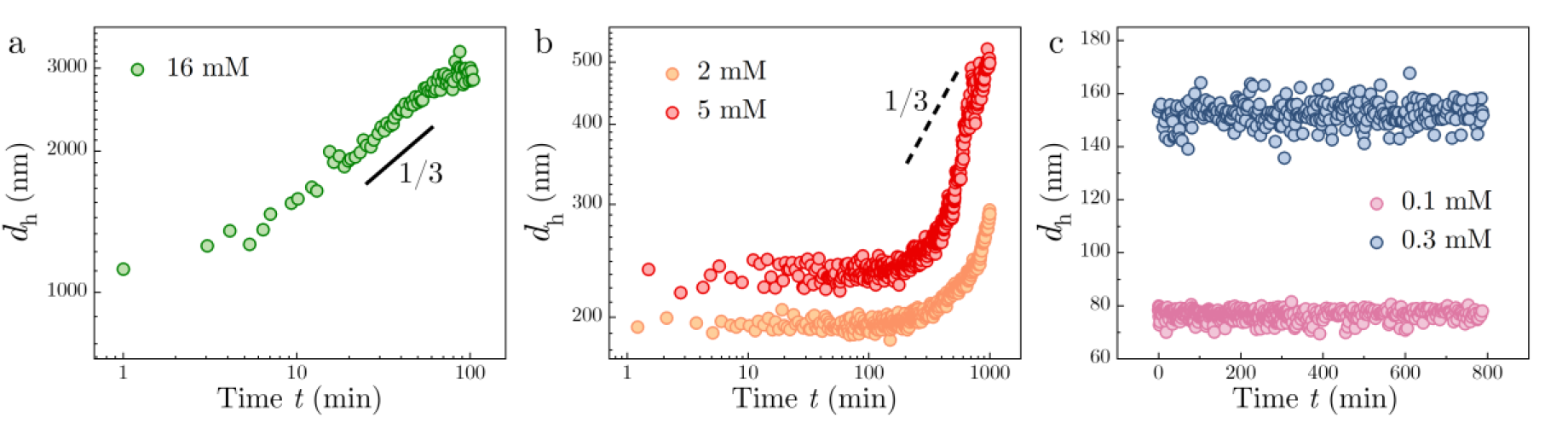
Nanoscale complex coacervates formed at low concentrations exhibit suppressed coarsening behavior. (a) For coacervates formed at *c*_m_ = 16 mM, the hydrodynamic diameter *d*_h_ exhibits a power-law increase with time, following the relationship *d*_h_~*t*^*α*^, where the exponent *α* ≈ 1/3. (b) Coacervates formed at *c*_m_ = 2 mM and *c*_m_ = 5 mM are initially stable but their *d*_h_ increase abruptly at later times with power-law exponents seemingly larger than 1/3. (c) Coacervates formed at *c*_m_ = 0.1 mM and *c*_m_ = 0.3 mM remain size-stable over 12 hours, demonstrating complete arrest of coarsening i.e., their mean hydrodynamic diameters remain constant throughout the observation period.

To demonstrate the universality of these anomalous coarsening behaviors, we repeat experiments using a different complex coacervate system: poly(vinylsulfonicacid, sodium salt) (PVS) and poly(diallyldimethylammonium chloride) (PDDA) mixed at a neutral charge ratio. Similarly, DLS reveals the formation of complex coacervates with *d*_h0_ ranging from tens of nanometers to a few microns (Supplementary Fig. 3, Supplementary Fig. 4, and Supplementary Fig. 5). The aforementioned three distinct types of coarsening behaviors are also observed for PVS/PDDA coacervates with different initial hydrodynamic diameters (Supplementary Fig. 6). Beyong synthetic coacervates, suppressed coarsening has recently been identified for various protein condensates, which exhibit power-law exponents smaller than 1/3 ^14,20–24^. In particular, some studies also echo that stable nanoscale condensates form more readily at low macromolecular concentrations ^21,23^. These results reveal three distinct coarsening regimes depending on concentration: (i) classical power-law growth, (ii) delayed growth, and (iii) long-term arrested growth. These collectively suggest a general mechanism underlying the suppressed coarsening.

### Size-dependent coarsening rate of nanoscale coacervates

To understand the mechanism that underlies these unexpected results, we start by reviewing two basic coarsening mechanisms: Brownian motion-induced coalescence (BMC) and Ostwald ripening. BMC is caused by coalescence of droplets that undergo stochastic Brownian motion ^1–3^, while Ostwald ripening is driven by flux of materials from smaller droplets to larger droplets due to the difference in their internal pressures and chemical potentials ^1,4,5^. Theoretically, both mechanisms follow the power-law scaling, 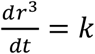, where *r* is the mean radius of droplets, *t* is the duration time, and *k* is a growth rate coefficient. However, the rate coefficients of BMC (*k*_BMC_) and Ostwald ripening (*k*_OR_) depend on distinct physical parameters (Supplementary Note 1). Given physical parameters of complex coacervates in this work, we find that *k*_BMC_ ≳ 3.9 × 10^−6^ μm^3^/s and *k*_OR_ ≈ 1.1 × 10^−7^ μm^3^/s (Supplementary Note 1). This suggests that BMC dominates coarsening, consistent with previous studies showing Ostwald ripening is generally negligible in coacervate systems ^1,16^. Therefore, we mainly focus on BMC as the dominant mechanism in the following sections. As a result, the coarsening behavior of microscale coacervates formed at *c*_m_ = 16 mM (Fig. 2a) can be described by the BMC mechanism, not only does the power-law exponent closely match 1/3, but the rate coefficient *k* also aligns with the theoretical estimate *k*_BMC_ (Supplementary Note 1). However, the coarsening behavior of nanoscale coacervates formed at low polyelectrolyte concentrations deviates significantly from the curves predicted by the BMC mechanism (Supplementary Fig. 7). Notably, prior work by Shimizu and Tanaka have proposed that in BMC scenarios, droplet motion can even be enhanced by hydrodynamic interactions such as Marangoni flows, which arise from interfacial tension gradients induced by compositional correlations among droplets [47].

Why is the coarsening of nanoscale coacervates suppressed? To answer this, we first calculate the growth rate 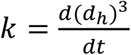 as a function of coacervate size for nanoscale coacervates growing at *c*_m_ = 2 mM (Supplementary Fig. 8). As shown in Fig. 3a, *k* depends on the coacervate size *d*_*h*_ and increases tremendously with *d*_*h*_ (Fig. 3a). Since coacervate droplets grow primarily via calescence in BMC, the results suggest that smaller coacervates merge much slower than larger ones. This steep increase in *k* over *d*_*h*_ can be described using a phenomenological exponential expression, *k* = *k*_0_*exp*[(*d*_*h*_/*d*_*c*_)^*n*^], where *k*_0_, *d*_*c*_, *n*, are fitted constants (Fig. 3a, red dashed line). In contrast, a power-law expression, defined as *k* = *k*_0_ + *a*(*d*_*h*_/*d*_*c*_)^*n*^, gives rise to a slow transition and fails to capture the experimental results.

**Fig. 3.**
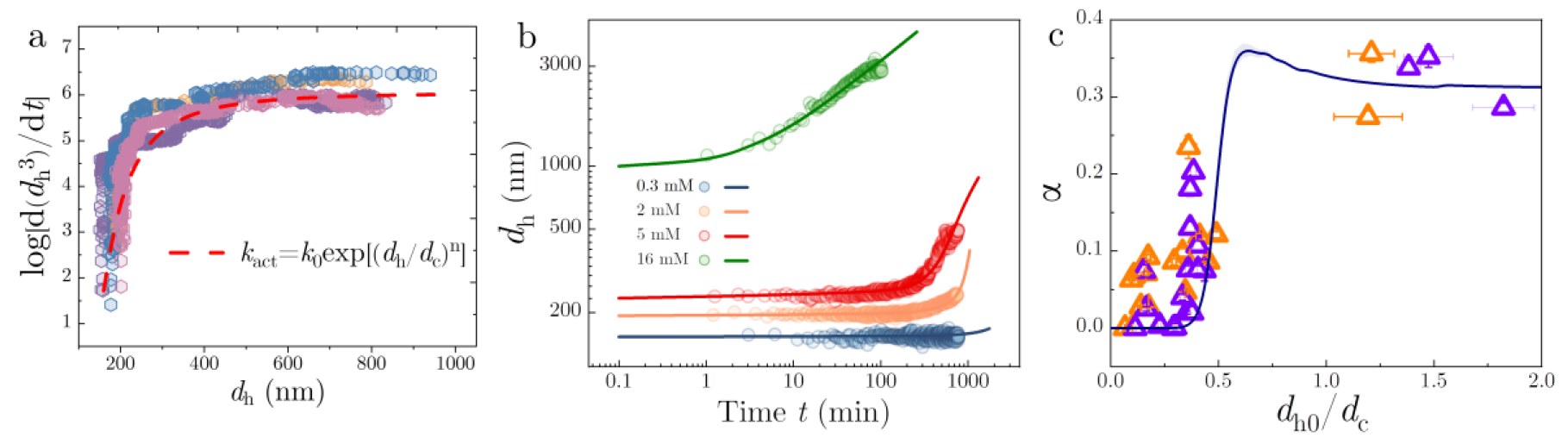
Merging-limited coarsening of nanoscale coacervates. (a) Plot of the logarithmic coarsening rate as a function of the coacervate diameter. Nanoscale coacervates are formed at a monomer concentration *c*_m_ = 2 mM and different colors represent different independent experimental trials. (b) The mean droplet hydrodynamic diameter *d*_*h*_ as a function of time, with experimental data represented by hollow circles and the theoretical results from Eq. (1) shown as solid lines. The parameters used in solving Eq. (1) are summarized in Supplementary Table 2. (c) The power-law exponent *α* of droplet coarsening during an initial time period (1-100 min) as a function of *d*_h0_/*d*_*c*_. The values of *α* are extracted from linear fitting of the experimental results on a log-log plot (PMA/PDDA coacervates: purple triangles; PVS/PDDA coacervates: orange triangles). The theoretical value of *α* (solid dark blue line) is obtained by fitting the theoretical curves of coarsening to a power law relation using Eq. (1) within the 1-100 min time period.

### Merging-limited coarsening governs the time evolution of nanoscale coacervates

To quantitively capture our experimental observations and elucidate the underlying mechanism, we develop a theoretical model that incorporates both droplet diffusion and size-dependent merging efficiency (see supplementary Note 2 for details) ^39^. The evolution of the droplet radius *r* is described by:

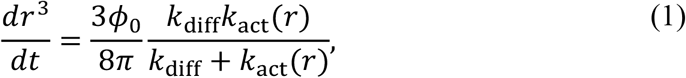

Here, *ϕ*_0_ is the droplet volume fraction, and *k*_diff_ = 16*πDr* is the encounter rate of diffusive droplets based on the Smoluchowski theory, where *D* is the diffusion constant of a droplet of radius *r* ^40^. The term *k*_act_ denotes the size dependent merging efficiency ^41^, quantifying the probability that two colliding droplets successfully coalesce. Based on the fitting in Fig. 3a, we adopt a phenomenological form for the merging rate as, 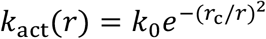, where *r*_*c*_ is a critical size that sets the merging barrier. In the limit where merging is much faster than diffusion, i.e., *k*_act_ ≫ *k*_diff_, Eq. (1) reduces to the classic BMC model ^3^,

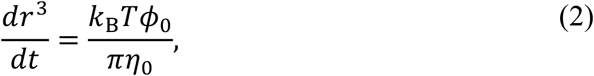

which leads to *r*~*t*^1/3^. In the opposite limit, where merging is extremely slow *k*_act_ ≪ *k*_diff_, we derive a merging-limited coarsening (MLC) model:

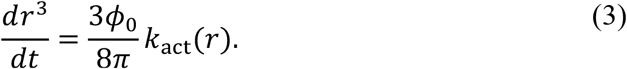

In the limit where *k*_act_ → 0, Eq. (3) simplifies to log *r* ~*k*_act_*t*, implying an exponential growth of *r* over time (Supplementary Note 2). To validate this, we replot experimental coarsening data of PMA/PDDA and PVS/PDDA coacervates within semi-logarithmic coordinates. Results show that nanoscale coacervates indeed exhibit linear growth of *d*_*h*_ over time in these plots, confirming an exponential growth via the MLC mechanism (Supplementary Fig. 9a and Supplementary Fig. 9c). By contrast, microscale coacervates that coarsen via BMC do not exhibit such linear growth in semi-logarithmic plots (Supplementary Fig. 9b and Supplementary Fig. 9d).

Fitting experimental data with Eq. (1) yields theoretical curves that quantitatively capture the coarsening behaviours of coacervates across varying initial sizes (Fig. 3b). In particular, it captures the anomalous growth at intermediate concentrations, where coacervates undergo a transition between MLC to BMC. By extracting a pseudo-exponent from fits of experimental results to the hypothesized power-law function *r*~*t*^*α*^ during early stages (~ 2 hours), we find that the exponent is smaller than 1/3 for small coacervates and plateaus at ~1/3 for coacervates with larger initial sizes (Supplementary Fig. 10). Such transition of *α* is also described by a theoretical curve that presents a rapid increase in *α* over *d*_h0_/*d*_*c*_ and a plateau at 1/3 when *d*_h0_/*d*_*c*_ > 1 (Fig. 3c). These observations align with recent experimental studies showing slow coarsening in condensates or coacervates with *α* < 1/3 ^22,42^. To validate our theoretical model, we also perform Monte Carlo simulations to replicate the coarsening process of coacervate droplets moving randomly by Brownian motions but with size-dependent merging efficiency (See Methods for details). The simulations incorporate a phenomenological relationship similar to that used in the analytic model to account for a size-dependent merging efficiency. The simulations results mirror experimental and theoretical observations: droplets grow slowly when small but exhibit BMC growth when large (Supplementary Fig. 11).

Similar to our observations with complex coacervates, a recent study show that condensates formed through the simple coacervation of intrinsically disordered proteins (IDPs), such as hnRNPA2 LCD, resist to grow under various conditions ^22^. To test the generality of our framework, we apply our theoretical model to analyze the coarsening data from this study ^22^. The BMC model effectively captures the rapid coarsening dynamics of condensates (Supplementary Fig. 12, red points), but deviates significantly when describing condensates with low growth rates (Supplementary Fig. 12, black points). In contrast, our proposed MLC model successfully captures these slow coarsening dynamics (Supplementary Fig. 12). These results highlight the broad applicability and physical relevance of the MLC mechanism across diverse condensate systems.

### Electrostatic repulsion leads to restricted merging of coacervate droplets

Having demonstrated the merging-limited coarsening, we then discuss the potential mechanisms that may reduce the merging efficiency between small-sized coacervate droplets. The MLC identified in this study shares similarities with the merging-limited aggregation in colloidal particles, where the colloidal aggregation occurs very slowly following an exponential function over time ^43^. In such colloidal systems, surface charge-induced electrostatic repulsions are widely recognized to reduce the aggregation and stabilize colloidal particles. Similarly, condensates or coacervates formed by charged macromolecules experience electrostatic repulsions that contribute to their size stability ^20,44^. These observations suggest that electrostatic interactions play a central role in suppressing droplet merging at the nanoscale.

To test the effect of electrostatic repulsion, we measure zeta potentials of PMA/PDDA coacervates. We find that these coacervates exhibit significant surface charge, while zeta potential slightly decreasing as concentration increases (Fig. 4a). For example, nanoscale coacervates formed at *c*_m_ = 2 mM exhibit a high negative zeta potential of approximately −20 mV (Fig. 4a). By measuring the zeta potential of coacervates over time, it shows that, as the droplet size increases, the absolute value of zeta potential decreases following a power-law relationship with a fitted exponent of ~0.37 (Supplementary Fig. 13). Coacervates formed at *c*_m_ = 5 mM exhibit a similar power-law decay, though with a smaller exponent of ~ 0.21 (Supplementary Fig. 13).

**Fig. 4.**
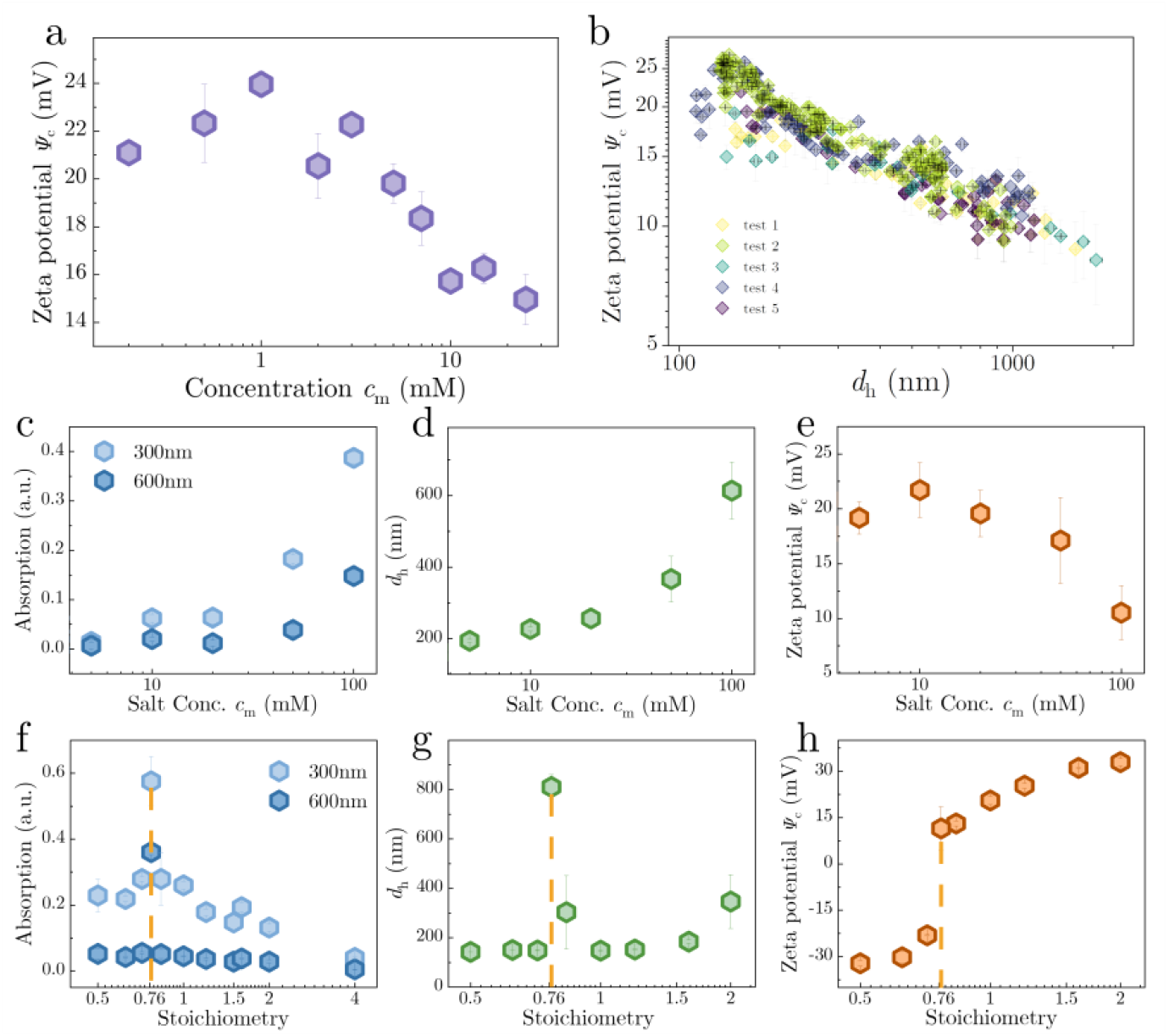
Charged coacervate interfaces underlies their stability and suppressed coarsening behavior. (a) Plot of the zeta potential of complex coacervates as a function of the monomer concentration *c*_m_. (b) Logarithmic plot of Zeta potential *Ψ*_*c*_ of coacervates as a function of their size. *Ψ*_*c*_ exhibits a power-law decay over size. Diamond symbols in different colors represent different independent tests with coacervates formed at *c*_m_ = 2 mM. Plots showing changes in solution absorption value (c), hydrodynamic diameter (d), and Zeta potential *Ψ*_*c*_ (e) of coacervates (*c*_m_ = 2 mM) with the addition of salts at different concentrations. Plots showing changes in solution absorption value (f), hydrodynamic diameter (g), and Zeta potential *Ψ*_*c*_ of coacervates (h) (*c*_m_ = 2 mM) at different stoichiometric ratio of PDDA to PMA. A neutral charged state of coacervates is observed when the stoichiometric ratio is approximately 1.3/1.

To further verify the effect of electrostatic repulsion, we add salt into the solution to screen electrostatic interactions. As anticipated, increasing salt concentration increases coacervate solution turbidity, evident both in spectroscopy measurement and bulk solutions in microcentrifugation tubes (Fig. 4c and Supplementary Fig. 14). Concomitantly, we observe that higher salt concentrations lead to larger initial coacervate sizes (Fig. 4d) and a decrease in the absolute value of zeta potential (Fig. 4e). Notably, at salt concentration higher than 50 mM, microscale coacervates become visible under an optical microscope (Supplementary Fig. 15). Moreover, coacervates formed at *c*_s_=50 mM start growing rapidly through BMC with an power-law exponent close to 1/3 (Supplementary Fig. 16). These results provide strong evidence that electrostatic repulsion among coacervate droplets underlies their long-term size stability.

### Chain length asymmetry-induced charge separation and electrostatic repulsion

While previous studies have shown that condensates or coacervates are charged, strong electrostatic potentials in these systems arise from imbalanced stoichiometric ratios of oppositely charged polymers ^45,46^. In that case, excessive portion of charged polymers will attach to condensate surfaces and generate an electrostatic potential. However, in our study, coacervates are all formed *under conditions of neutral stoichiometry*, which in principle promotes efficient phase separation and minimize net charge within condensates. This then raises a critical question: what causes the strong surface charge observed in coacervates formed at neutral stoichiometry?

To answer this question, it is noted that the polyelectrolytes used to form our coacervates have unequal chain lengths, with PDDA chains being substantially longer than those of PMA. Therefore, these results suggest the hypothesis that chain-length asymmetry promotes entropic partitioning, whereby shorter chains preferentially partition into the dilute phase to maximize their configurational entropy. This asymmetric distribution induces interfacial charge imbalance even at globally neutral stoichiometry, leading to electrostatic repulsion that stabilizes the coacervates. To test whether chain-length asymmetry contributes to the observed surface potential and stability of nanoscale condensates, we systematically varied the stoichiometric ratio of charged monomers (PDDA to PMA) to tune the surface charge of the coacervates. When the ratio exceeds 1— indicating an excess of PDDA—coacervates retain high surface charge density and exhibit pronounced stability (Fig. 4h). In contrast, at ratios below 1, the coacervates become less stable, as evidenced by larger initial sizes, increased turbidity, and reduced zeta potentials (Fig. 4f–h). Notably, a distinct peak in instability emerges near a stoichiometric ratio of ~1/1.3, suggesting that coacervates are most unstable under this condition (Fig. 4f and 4g). These findings imply a strong entropic contribution, where chain-length asymmetry and dilution lead to asymmetric partitioning and interfacial charge separation:

To further validate the role of chain-length asymmetry, we perform fluid particle dynamic (FPD) simulations ^47–49^, that incorporates both many-body hydrodynamic and long-range electrostatic interactions (see Methods for details of FPD simulations). We begin with an asymmetric chain length configuration in which each positively charged chain consists of 100 monomers, whereas each negatively charged polymer chain contains 10 monomers. All other parameters, such as monomer concentration, size, and valency, were kept constant. To preserve overall charge neutrality, simulations include a greater number of negatively charged chains than positively charged ones.

Initial simulations at a monomer concentration of 92.5 mM show power-law growth with an exponent approaching 1/3, consistent with the conventional BMC mechanism (Fig. 5a and Fig. 5c, yellow points). However, when the monomer concentration is reduced to 27.4 mM, the coacervates exhibit slower coarsening, particularly during the late stage with an exponent smaller than 1/3 (Fig. 5a and Fig. 5c, blue points). Although the early-stage growth superficially resembles classical Brownian motion-induced coalescence (BMC), it may not strictly follow the standard BMC mechanism. Prior work has shown that droplet motion can be driven by Marangoni forces arising from interfacial tension gradients induced by compositional correlations between droplets ^47^. However, such hydrodynamic effects likely become irrelevant for nanoscale droplets due to insufficient timescale separation between droplet and polymer diffusion, precluding the establishment of quasi-equilibrium concentration fields.

**Fig. 5.**
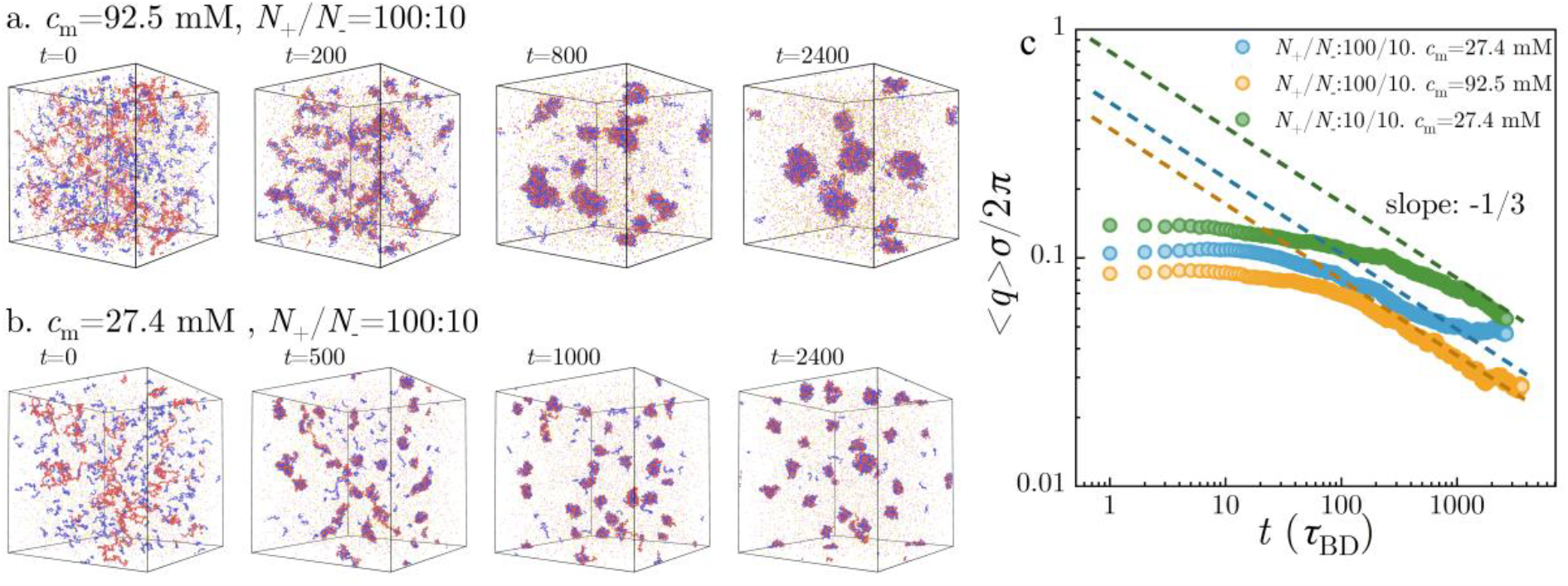
The suppressed coarsening behavior of coacervates is attributed to the combination of asymmetric chain length and low molecular concentration. Structural evolution of simulated coacervates with asymmetric chain lengths at different time with a unit of *τ*_BD_ that is the Brownian time of a monomer under Bjerrum length *l*_B_=0.72nm (~1.115σ σ is the diameter of a monomer). (a) *c*_m_=92.5 mM and *N*_+_/*N*_−_=100:10. (b) *c*_m_ =27.4 mM and *N*_+_/*N*_−_=100:10. Red and blue beads represent monomers of polycations and polyanions. Counterions are depicted as smaller particles with magenta and yellow beads indicating associated counterions with polycations and polyanions, respectively. All simulations contain 25 polycations and 250 polyanions. (c) Temporal evolution of characteristic wavenumber <*q*>, defined as the first moment of structural factor *S*<*q, t*> in FPD simulations (see Methods for details of <*q*>). Simulations include two asymmetric cases: *c*_m_=92.5 mM and *N*_+_/*N*_−_=100:10; *c*_m_ =27.4 mM and *N*_+_/*N*_−_=100:10 and one symmetric case: *c*_m_ =27.4 mM and *N*_+_/*N*_−_=10:10. The observed plateau in ⟨q⟩ at late stages may correspond to the experimentally observed arrest of coarsening, consistent with a scenario in which entropic charge separation induces electrostatic repulsion that suppresses merging.

We propose that entropic effects arising from chain-length asymmetry introduce interfacial charge separation, thereby generating size-dependent electrostatic barriers. This mechanism operates even under stoichiometrically balanced conditions, offering a unifying explanation for suppressed coarsening across diverse systems. The resulting associative complexation imposes microscopic kinetic constraints that reshape the coarsening pathway into a two-step process: (1) rapid formation of ionically bound complexes and (2) their mesoscale growth and coalescence. Consequently, the early-stage growth captured in our FPD simulations may reflect this non-classical pathway rather than conventional BMC.

These two comparison simulations align closely with experimental observations, indicating that coarsening suppression is exclusive to relatively low monomer concentrations. To prove the necessity of chain length asymmetry, we perform an additional simulation in which both positively and negatively charged polymer chains comprise 10 monomers. As a result, the simulation shows a typical rapid coarsening with an exponent close to 1/3 (Fig. 5c, green points, and supplementary Fig. 17). This confirms that chain length asymmetry is critical for inducing suppressed coarsening.

As shown in Fig. 6a–c, coacervates formed with asymmetric chain lengths and low concentrations exhibit a higher density of unsaturated positively charged residues at the interface, along with a larger population of short polyanions dispersed in the dilute phase. This reflects entropic partitioning driven by chain-length asymmetry, where shorter chains favor the dilute phase to maximize their configurational entropy—leading to interfacial charge separation even under globally neutral stoichiometry. Analysis of coacervate charge densities reveals that most coacervates formed with asymmetric chain lengths at low concentration carry more positive charges (Fig. 6d). However, as concentration increases, a subset of coacervates acquires negative charge density, reducing the size-averaged charge density (Fig. 6b, Fig. 6d and supplementary Fig. 18). This trend aligns with zeta potential measurements over concentration in Fig. 4a. In both concentrations, charge density and net charge of coacervates decreases monotonically with their size (Fig. 6d and Supplementary Fig. 19), consistent with zeta potential measurements over size (Fig. 4b). In contrast, coacervates formed with symmetric chain lengths exhibit near-zero net charge density regardless of size (Fig. 6d). These results underscore the critical role of chain length asymmetry in activating the MLC mechanism.

**Fig. 6.**
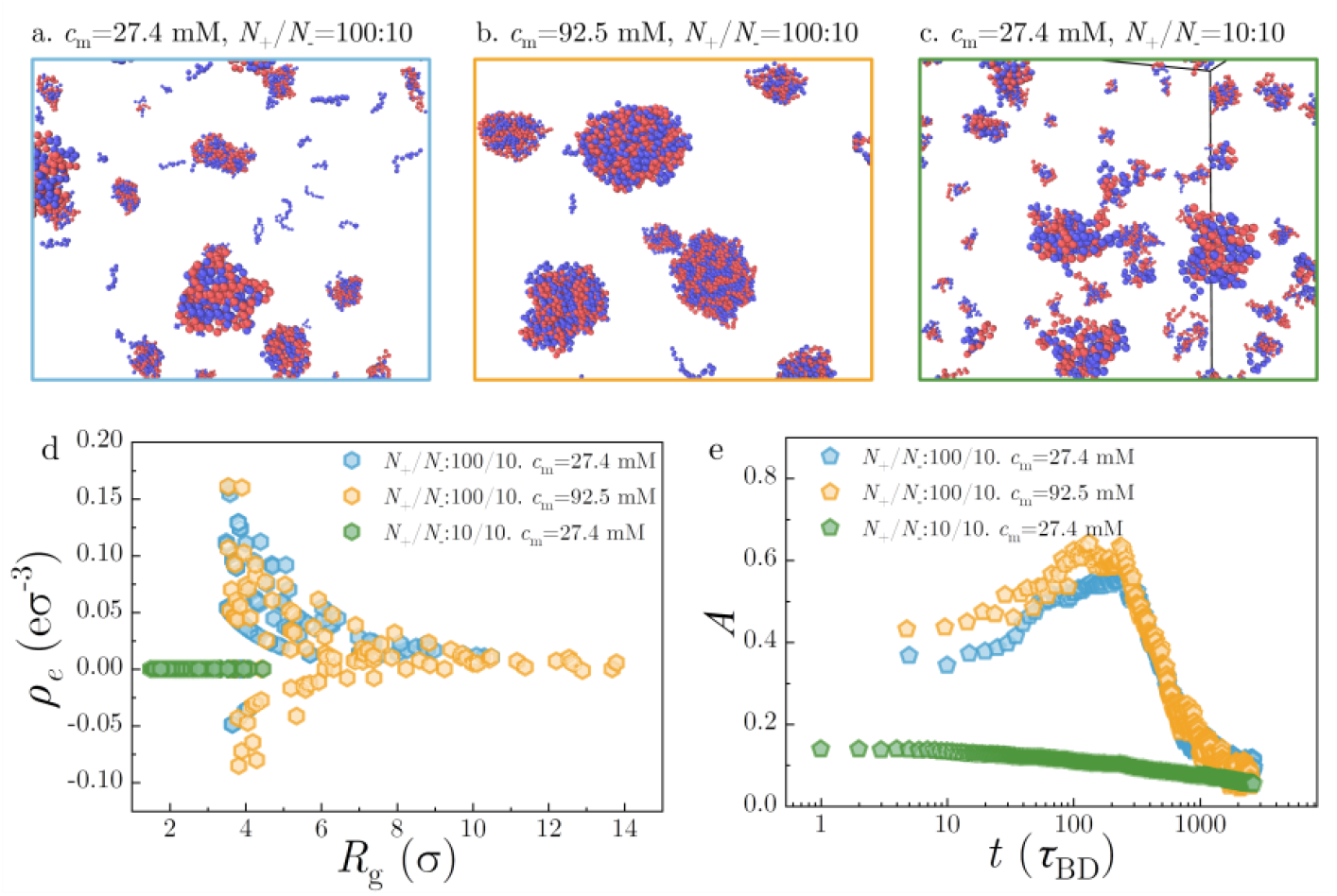
Asymmetric chain lengths induce charge separation at interface and electrostatic repulsions among complex coacervates. Representative structural configurations of coacervates formed under conditions of (a) *c*_m_=27.4 mM and *N*_+_/*N*_−_=100:10, (b) *c*_m_ =92.5 mM and *N*_+_/*N*_−_=100:10, and (c) *c*_m_ =27.4 mM and *N*_+_/*N*_−_ =10:10. Freely dispersed polyanion chains are observed mostly in the case (a). (d) Plot of the charge density of individual droplets as a function of their size with the unit of σ. (e) Temporal evolution of the degree of asphericity *A* of individual droplets over time.

Moreover, analysis of the asphericity *A* of coacervates shows that coacervates with polyelectrolytes of asymmetric chain lengths adopt non-spherical morphologies (Fig. 6e), including “tadpole”-like and “fold”-like shapes, during the early growth phase (Fig. 6a). These deviations from spherical globules arise from chain length asymmetry and weak shrinking tendency of nanoscale droplets^49^, which induces the formation of loosely packed, extended structures with net charges. Analogous cluster geometries have also been observed in symmetric polycation/polyanion coacervates when the stoichiometry is not neutral ^45,46^. Overall, the FPD simulations qualitatively capture our experimental observations and clarify the molecular mechanisms underlying the chain length asymmetry-induced merging limited coarsening of coacervates.

## Discussions

In this work, we combined experiments, theory, and simulations to uncover a novel mechanism underlying the suppressed coarsening in complex coacervates or biomolecular condensates. The proposed mechanism, in which size-dependent electrostatic barriers strongly hinder droplet merging, enables extremely slow coarsening kinetics and long-term size stability of nanoscale condensates. In contrast to conventional LLPS that is purely driven by free energy minimization, the formation of nanoscale complex coacervates begins with molecular-level interactions, such as electrostatic interactions between oppositely charged polymers. These initial interactions impose microscopic kinetic constraints, setting the stage for unique coarsening behaviors that lead to distinct features in nanoscale coacervates, such as sensitivity to chain length asymmetry, entropic charge separation, and thus size-dependent electrostatic barriers that underlies the suppressed coarsening behaviors.

A conceptually related phenomenon is the moving droplet phase (MDP) observed in viscoelastic phase separation, where droplets resist coalescence by behaving like elastic particles upon collision ^50^. However, MDP relies on fast droplet motion and slow internal viscoelastic relaxation—conditions unlikely to arise in crowded intracellular environments. More critically, MDP does not account for the pronounced salt dependence of droplet merging observed in our system, which points to an electrostatic rather than mechanical origin. Notably, unlike MDP, which imposes a rheology-dependent, dynamic barrier to coalescence, the merging-limited coarsening (MLC) mechanism introduces a static, size-dependent electrostatic barrier, offering a more general and robust framework for stabilizing biological condensates.

Classical coarsening models of LLPS emphasize thermodynamic free energy minimization, typically predicting continuous, unimpeded droplet coarsening. This traditional view treats the growth of condensates with symmetric chains and balanced interfacial electrostatics as barrier-free and thermodynamically downhill (Fig. 7c, d). In contrast, our results highlight the critical role of kinetic constraints—particularly electrostatic merging barriers—arising from interfacial charge separation. These barriers create size-dependent merging inefficiencies that arrest coarsening at the nanoscale. The MLC framework challenges the classical LLPS framework, highlighting that microscopic constraints, such as chain-length asymmetry and interfacial electrostatics, can impose kinetic bottlenecks that dominate droplet dynamics. Consequently, LLPS proceeds not along a smooth thermodynamic descent but over a rugged energy landscape shaped by molecular interactions and interfacial physics (Fig. 7a, b). These droplets are kinetically trapped and can only grow by overcoming merging barriers, resulting in markedly slowed coarsening.

**Fig. 7.**
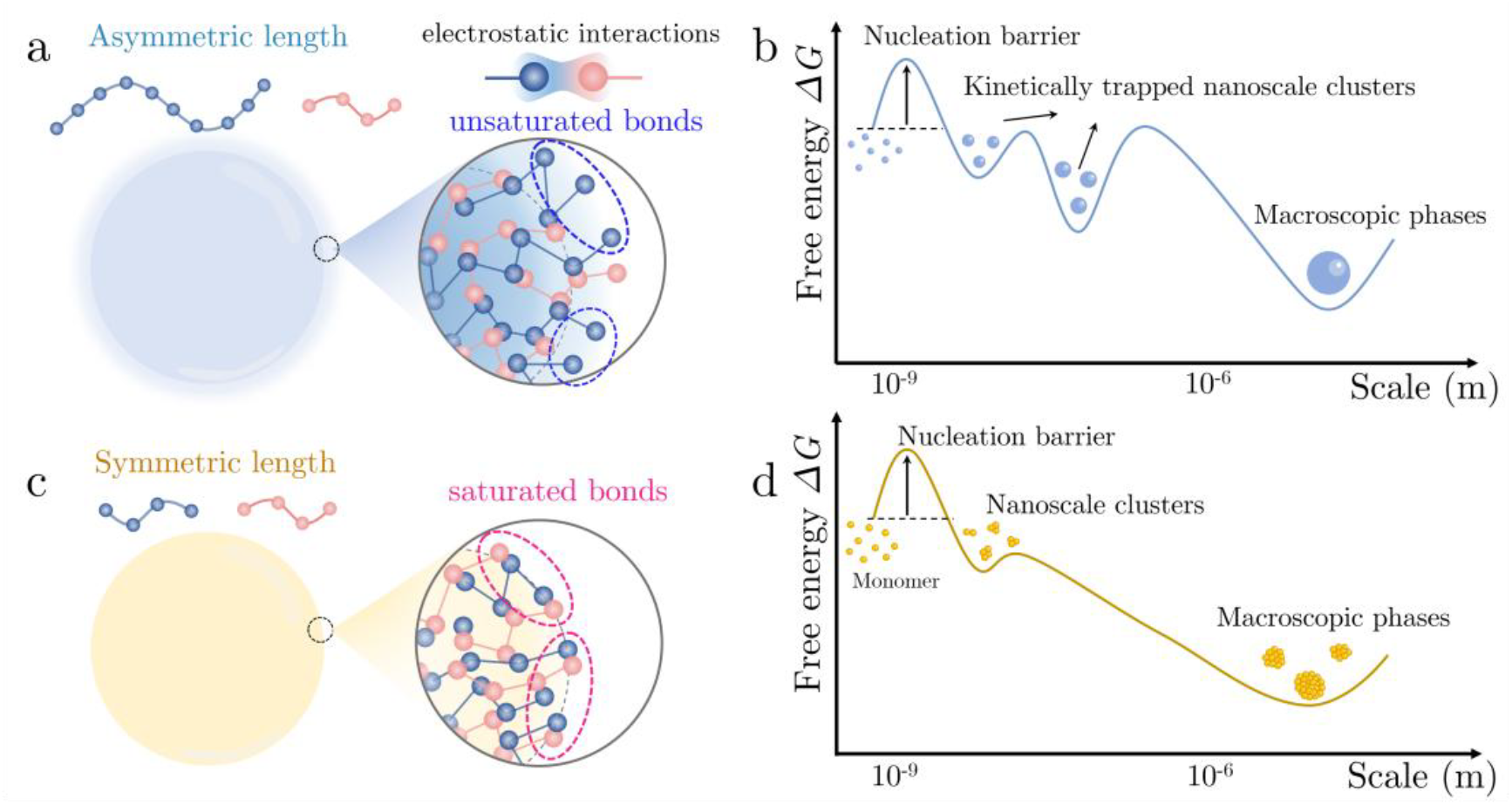
Nanoscale coacervates featuring asymmetric chain exhibit a rugged energy landscape of coarsening. (a)(b) Schematic illustrations of coacervates with asymmetric chain lengths that have unsaturated charged residues at the interface and their coarsening process is represented as a rugged terrain comprised of multiple local energy minimum separated by mountainous energy barriers. (c)(d) In contrast, coacervates with symmetric chain lengths have neutralized interface, characterized by balanced electrostatic interactions at the interface. For these symmetric coacervates, the energy landscape associated with their coarsening is a smooth downhill path toward the global energy minimum.

This framework also accommodates other known stabilization mechanisms, including the formation of interfacial protein clusters ^12,51^ and surface charge accumulation ^44,52,53^, both of which lower interfacial tension and create physical impediments to merging. Moreover, the increasingly recognized role of interface resistance in reducing molecular transport across condensate boundaries offers another route to suppressed growth ^54,55^, further reinforcing the idea that condensate coarsening can be shaped by complex interfacial physics rather than being driven by thermodynamics alone.

Moreover, our results indicate that the stability of nanoscale condensates is most robust at relatively low concentrations. This low-concentration regime is physiologically relevant, as most phase-separated biomolecules in cells exist near their saturation concentrations (nanomolar to micromolar)^20^, which corresponds to the concentration window where we observe stable nanoscale droplets. These stable nanoscale condensates may serve critical functions in biology ^7,56^, including postsynaptic density in neurotransmission ^57^, transcriptional condensates in gene transcription ^26,58,59^, and heterochromatin in chromatin compaction ^60,61^. Therefore, our work may have provided mechanistic insights that explain the widespread size stability of condensates observed in living cells and in vitro reconstituted systems ^20,21,23,35,62^.

Our findings not only provide a mechanistic basis for condensate stability in biological contexts but also offer general design principles for engineering long-lived nanoscale assemblies. In biological contexts, cells may exploit these same physical principles to regulate condensate stability. By tuning effective chain length or macromolecular concentration, cells could modulate interfacial charge asymmetry and thus control whether condensates remain dynamic or enter metastable arrested states. Moreover, our findings offer a design principle for synthetic biology and materials science: by adjusting polyelectrolyte size asymmetry or concentration, one can engineer condensates with programmable lifetimes and sizes—enabling the creation of persistent or reconfigurable biomolecular assemblies. From a broader perspective, by revealing a non-classical coarsening mechanism governed by entropic electrostatics, our study expands the physical framework of phase separation and opens new avenues for controlling metastability in soft matter and nonequilibrium systems.

## Methods

### Preparation of complex coacervates

Stock solutions of poly(methacrylic acid, sodium salt) (PMA, Sigma-Aldrich, 674044), poly(vinylsulfonic acid, sodium salt) (PVS, Macklin, P909334), and Poly(diallyldimethylammonium chloride) (PDDA, Sigma-Aldrich, 522376) were firstly prepared at high concentrations using Milli-Q water (18.2 MΩ, pH=7). The stock solutions were diluted according to designed *c*_m_. Complex coacervates were formed by gently mixing diluted oppositely charged polymer solutions at a charge-balanced neutral stoichiometry.

### Dynamic light scattering (DLS)

DLS measurements were preformed using a Zetasizer Pro (Malvern Instruments, UK) with a measurement range from 0.3 nm to 10 μm. Complex coacervate samples with volume of approximately 1mL were transferred into transparent cuvettes (DTS0012, Malvern) using a pipette. The cuvettes were subsequently sealed with a plastic cap and placed into the Zetasizer Pro machine. A He-Ne laser (633 nm) was used to illuminate the sample and the intensity of light scattered at a constant angle of 173° was measured using a photodiode. The program presets the viscosity of water (1.0 mPa) and the refractive index (1.38) for the measurement, settled at room temperature (25 °C) with a 120 s equilibration time. To determine the hydrodynamic diameters *d*_*h*0_, three measurements were carried out for each sample, and at least three independent samples were measured to obtain a mean ± SD. To conduct the long-term monitoring of *d*_*h*_, the program was set to continuously record the hydrodynamic diameters at intervals ranging from 1 to 3 minutes.

### Optical microscopy

The bright-field images were captured by an inverted fluorescence microscope (Olympus) equipped with a differential interference contrast (DIC) filter. The sample was loaded into customized chambers fabricated using spacers (ThermoFisher, S24737) sandwiched by two coverslides at a 120 μm gap. The coverslides were cleaned by ethanol and water to remove contaminants and coated with mPEG-Silane (Aladdin, S164281) to reduce the adhesion of coacervates onto surfaces. The coated slides were carefully rinsed using Milli-Q water to remove chemical residues, followed by drying inside 65 °C oven for 1 hour.

### Transmission electron microscopy (TEM)

Carbon-coated copper grids were firstly discharged by a Glow Discharge Cleaning System to become hydrophilic (Pelco EasiGlow 91000). 5 μL samples containing complex coacervates was loaded onto the grids (Ted Pella, Inc.). The samples on grids were negatively stained with uranyl acetate (UA), followed by general removal of excess staining solutions using a filter paper wick, and subsequent air-drying before imaging. The TEM were performed using a FEI Tecnai G2 20 facility at the Electron Microscope Unit (EMU) at the University of Hong Kong.

### Monte Carlo Simulations

The Monte-Carlo simulations were performed in MATLAB. Initially, a total of *N* droplets were generated in a three-dimensional box of dimensions 35×35×35 using random seeds, with periodic boundary conditions. At each time step, droplets were allowed to move randomly in three dimensions at a distance set by the mean square displacement (MSD), that is ⟨*x*⟩^2^ = 6*D*Δ*t* where the diffusion coefficient was determined by the Stoke-Einstein equations *D* = *k*_*B*_*T*/(6*πηr*). In the case of BMC, if droplets overlapped, they were merged to form a single large droplet located at their center of mass. The size of new droplet was determined by the volume and mass conservation of two overlapped droplets. In our modified model, in the case of overlapping droplets, the merging efficiency *E* was calculated by Eq. (2) using the mean radius of two overlapped droplets. A random number was generated as *P* for each merging counter. If *P*≤ *E*, the two droplets were merged as previously described. On the other hand, if *P* > *E*, the droplets would move away from each other in opposite directions parallel to the line connecting their centers. The mean radius *r* of all droplets was calculated and recorded at each timepoint.

### Simulation Model of polyelectrolytes

We employ a coarse-grained polyelectrolyte model in which monomers are modeled as spherical monomers of diameter σ and connected by harmonic bonds to maintain the chain connectivity. The system contains positively charged polycations and negatively charged polyanions, along with monovalent counterions for charge neutrality. The ions and monomers are placed in a cubic periodic simulation box and interact via a purely repulsive shifted-truncated Lennard-Jones (LJ) potential with a coupling constant *ε*_*LJ*_ = *k*_*B*_*T*, where *k*_*B*_ is Boltzmann’s constant and *T* is the absolute temperature. As customary for polyelectrolyte simulations, we adopt a Bjerrum length *l*_*B*_ = 0.72 nm ~1.1 σ. The covalent bonds between the neighboring monomers are modelled by a harmonic potential *V*_*bond*_ = (*K*/2)(*r* − *r*_0_)^2^ with a spring constant *K* = 400*k*_*B*_*T*/ σ^2^ and an equilibrium length *r*_0_ = 2^1/6^σ. We first use purely repulsive LJ interactions to establish a homogeneous initial state. Phase separation is then triggered by activating electrostatic interactions, which is computed via Ewald summation with a relative force accuracy of 10^−3^.

### Fluid particle dynamics

It is well known that hydrodynamic interaction (HI) is essential for properly modelling the non-equilibrium phase separation dynamics. To incorporate HI, we use the fluid particle dynamics (FPD) method ^47–49^. This approach, originally developed by Tanaka ^47^, has recently been applied to study the conformational transition of a polyelectrolyte chain ^48^ and the network-forming phase separation of oppositely charged polyelectrolytes ^49^. In the FPD framework, each particle n is represented as a viscous fluid particle described by a phase field ψ_*n*_(*r*) = 1/2{*tanh* [(*a*−| ***r*** − ***R***_***n***_ |)/*ξ*] + 1}where **r** is the position vector of the field, a is the particle radius, *ξ* is the interfacial thickness, and **R**_n_ is the position vector of particle n. This formulation allows us to construct a spatially varying viscosity field *η*(*r*) = *η*_*s*_ + (*η*_*p*_ − *η*_*s*_)∑_*n*_ψ_*n*_(*r*) where *η*_*s*_ and *η*_*p*_ are the solvent and particle viscosities, respectively. The flow field ***ν*** is obtained by solving the Navier-Stokes (NS) equation *ρ*(*∂*/*∂t* + ***ν*** *·∇*)***ν*** = ***f*** +*∇ ·* (***σ*** + ***σ***^***R***^) where *ρ* is the fluid density, ***f*** = ∑_*n*_***f***_***n***_ψ_*n*_(***r***) / ***∫***ψ_*n*_(***r***)*d****r*** is the force field derived from the smeared-out interaction forces ***f***_***n***_ acting on the particle n (including LJ interaction force and electrostatic force), and ***σ*** = *η*(***r***)[*∇****ν*** + (*∇****ν***^*^)***T***] −***PI***is the viscous stress tensor with ***I***the unit tensor and p the pressure enforcing incompressibility *∇*·***ν*** = 0. The term ***σ***^***R***^ is the random stress field, which satisfies the fluctuation-dissipation theorem 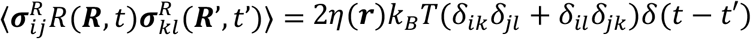 where *δ*(·) denotes the Dirac delta function. We solve the NS equation using the Marker-and-Cell (MAC) method on a staggered grid under three-dimensional periodic boundary conditions, ensuring accurate resolution of the solvent flow. Electrostatic interactions are calculated using the Ewald summation with a relative force accuracy of 10^−4^. The FPD simulations were carried out using NVIDIA A40 GPU.

### Structure factor *S*(*q,t*) and characteristic wavenumber ⟨*q*⟩

To quantify coarsening dynamics in FPD simulations, we compute the structure factor *S*(***q***, *t*) from the three-dimensional power spectrum of the coarse-grained density field *S*(***q***, *t*) = *ρ*_***q***_(*t*)*ρ*_−***q***_(*t*)/*N*, where *ρ*_***q***_(*t*) is the Fourier transform of the density field *ρ*(***r***, *t*) defined as *ρ*(***r***, *t*) = ∑_*n*_ψ_*n*_(*a*−| ***r*** − ***R***_*n*_(*t*) |). Here, ψ_n_ is the hyperbolic tangent function representing the phase field of the n-th particle as in the FPD method. The characteristic wavenumber ⟨***q***⟩, which reflects the characteristic length scale of the evolving phase separation domains, is obtained as ⟨*q*⟩ = ***∫****d*_*qq*_*S*(*q, t*)/***∫****d*_*q*_*S*(*q, t*).

### Charge density and degree of asphericity of droplets

In our FPD simulations, individual droplet clusters are defined based on the geometric criterion. Specifically, two monomers are considered to belong to the same coacervate droplet if the center-to-center distance between them is less than 1.5σwhere σis the particle diameter. The degree of asphericity A is measured to characterize the shapes of clusters. For each cluster, we construct the radius of gyration tensor 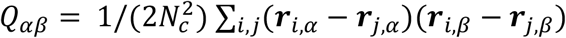, where ***r***_*i*_ is the position vector of the i-th monomer of the cluster, *α, β* = *x,y,z* are Cartesian components, and *N*_*c*_ is the number of monomers of the cluster. We obtain the radius of gyration *R*_*g*_ = (*λ*_1_ + *λ*_2_ + *λ*_3_)^1/2^ and the degree of asphericity *A*= ((*λ*_1_ − *λ*_2_)2 + (*λ*_2_ − *λ*_3_)2 + (*λ*_3_ − *λ*_1_)2)/(2(*λ*_1_ + *λ*_2_ + *λ*_3_)^2^) where*λ*_1_≤ *λ*_2_≤ *λ*_3_ are the eigenvalues of the radius of gyration tensor *Q*. The value of *A* is between 0 (sphere-like) and 1 (rod-like). The charge density *ρ*_*e*_of the individual droplets is measured as *ρ*_*e*_= *Q*_*e*_/(4/3*πR*_*g*_^3^).

## Acknowledgements

We thank Dr. Wei Guo and Dr. Xiufeng Li for useful discussions. This work was supported by the General Research Fund (Nos. 17306221, 17317322, and 17306820) from the Research Grants Council (RGC) of Hong Kong, as well as the National Natural Science Foundation of China (NSFC)-RGC Joint Research Scheme (N_HKU718/19). H.C.S. was funded in part by the RGC Senior Research Fellow (SRFS2425-7S04) from the RGC, the Croucher Senior Research Fellowship from Croucher Foundation and the Health@InnoHK program of the Innovation and Technology Commission of the Hong Kong SAR Government. H.T. was supported by the Grant-in-Aid for Specially Promoted Research (JSPS KAKENHI Grant No. JP20H05619) from the Japan Society for the Promotion of Science (JSPS). Y. Z. was supported by a startup fund at Johns Hopkins University. J. Y. was supported by the startup fund (Project Number: G0101000261-1003) provided by HKUST(GZ).

## Author contributions

F.C. conceptualized and designed the project. F.C. performed experiments, Monte Carlo simulations, and data analysis. F.C. and Y.Z. built the analytical models. J. Y. and H.T. performed fluid particle dynamics simulations and data analysis. J.Y., Y.Z., H.T., and H.C.S. supervised the project. F.C. wrote the manuscript, and all authors have revised it.

